# Maternal and paternal sugar consumption interact to modify offspring life history and physiology

**DOI:** 10.1101/2021.08.11.456016

**Authors:** Tara-Lyn Camilleri-Carter, Matthew D.W. Piper, Rebecca L. Robker, Damian K. Dowling

## Abstract

Intergenerational effects on offspring phenotypes occur in response to variation in both maternal and paternal nutrition. Because the combined maternal and paternal effects are rarely considered together however, their relative contributions, and the capacity for interactions between parental diets to shape offspring life history and physiology are not understood. To address this, we altered sucrose levels of adult fruit flies (*Drosophila melanogaster*) prior to mating, across two generations, producing parent-parent and parent-offspring combinations that were either matched or mismatched in dietary sucrose. We then measured lifespan, fecundity, body mass, and triglyceride levels in parents and offspring. We reveal complex non-additive interactions, that involve diets of each parent and offspring to shape offspring phenotypes, but the effects were generally not consistent with an adaptive response to parental diet. Notably, we find that interacting parental flies (sires and dams) lived longer when their sucrose treatments were matched, but they produced shorter-lived offspring.

## Introduction

Parents contribute to the development of their offspring beyond the direct genotypic effects of gene transfer (Bonduriansky, Crean, & Day, 2012; Gluckman, Hanson, & Low, 2019; Nystrand, Cassidy, & Dowling, 2016). Non-genetic parental effects can arise through either condition-dependant mechanisms (e.g. direct effects of variation in parental care), or through changes in the regulation of gene expression via environmentally-mediated epigenetic mechanisms (Curley, Mashoodh, & Champagne, 2017). Consequently, when the environment of a parent varies, this can affect parental contributions to their offspring, and shape offspring fitness (Marshall & Uller, 2007; Mousseau & Dingle, 1991; Mousseau & Fox, 1998; Mousseau, Uller, Wapstra, & Badyaev, 2009; Uller, Nakagawa, & English, 2013). These effects are plastic responses that occur across generations, termed intergenerational plasticity (when effects span one generation) and transgenerational plasticity (when effects span multiple generations). Intergenerational plasticity has been documented broadly— from bacteria, to fungi, to plants, and in both invertebrate and vertebrate animals (Dyer et al., 2010; Jablonka & Raz, 2009; Roach & Wulff, 1987). Such plasticity can be triggered in response to a wide range of parental environmental stresses or changes, such as challenges to immunity, nutrition, temperature, toxins, circadian rhythm, and light quality (Baker, Sultan, Lopez-Ichikawa, & Waterman, 2019; Bell & Hellmann, 2019; Donelan et al., 2020; Nystrand & Dowling, 2014; Sultan, Barton, & Wilczek, 2009).

Currently, two questions remain unresolved when it comes to understanding the broader implications and mechanisms underpinning environmentally-mediated (non-genetic) intergenerational plasticity. The first question is whether this plasticity is adaptive to offspring. Numerous studies have suggested that individuals exposed to particular environmental stresses can prime their offspring through mechanisms of non-genetic inheritance to cope with these same stresses (anticipatory parental effects), thereby augmenting offspring resilience and fitness (Marshall & Uller, 2007; Rowiński et al., 2020). Several meta-analyses however, have generally found evidence for anticipatory parental effects is limited (Radersma, Hegg, Noble, & Uller, 2018; Sánchez-Tójar et al., 2020; Uller et al., 2013; Yin, Zhou, Lin, Li, & Zhang, 2019). Progress in resolving this question has been somewhat hindered by a lack of experimental studies with the power to satisfactorily disentangle adaptive from non-adaptive intergenerational responses (Ivimey-Cook et al., 2020.; Sánchez-Tójar et al., 2020; Uller et al., 2013). These designs require both parents and their offspring are provided with both a control treatment and a novel environmental treatment, in all possible matched and mismatched combinations, thus enabling determination of whether offspring fitness is higher when offspring environment matches that of their parents (Burgess & Marshall, 2014; Sánchez-Tójar et al., 2020; Uller et al., 2013). Notwithstanding, even the use of such designs may be ineffective at partitioning out transfer from parent to offspring of condition-dependent effects, from transfer of adaptive anticipatory effects (Bonduriansky et al., 2012; Engqvist & Reinhold, 2016), and thus should be interpreted cautiously.

Second, the relative magnitude of paternal effects to maternal effects on offspring phenotypic expression remains ambiguous. While maternal effects have been studied for decades and are known to be pervasive (Mousseau & Fox, 1998), the possibility for non-genetic paternal effects to shape phenotypic expression in offspring received much less attention until recently (Crean & Bonduriansky, 2014; Immler, 2018). In the past decade however, it has become clear that males contribute to offspring phenotypes beyond that of their direct genotypic contributions (Crean & Bonduriansky, 2014; Evans, Wilson, Pilastro, & Garcia-Gonzalez, 2019; Immler, 2018). Despite recent progress, the relative contributions of paternal and maternal effects on offspring performance remain elusive, as does the question of whether paternal contributions interact non-additively with maternal contributions to shape intergenerational fitness in ways that may not be captured simply by measuring maternal or paternal contributions in isolation.

Dietary variation refers to heterogeneity across individuals in the quality or quantity of macronutrients ingested, and represents a major source of environmental influence in natural populations. Dietary variation affects a wide range of fitness related traits—from physiological measures of obesity, to reproductive success, and lifespan (Duxbury & Chapman, 2020). Many of these effects appear to be conserved across invertebrates and vertebrates, and modifications to diet in one generation have been shown to trigger indirect effects on the metabolic performance and body composition of offspring and grand offspring (Camilleri-Carter, Dowling, Robker, & Piper, 2019; Dunn & Bale, 2009; Ivimey-Cook et al., 2020). Research into dietary-mediated intergenerational inheritance has focussed on mice and flies, where studies have explored the intergenerational consequences of obesogenic diets. Recent insights in each system reveal persistent parentally-mediated effects of high fat (in mice) or high sugar (in flies) diets on offspring phenotypes, with effects that can transcend multiple generations (Buescher et al., 2013; Huypens et al., 2016; Öst et al., 2014). In particular, a recent study by Huypens et al. (2016) in mice showed that maternal and paternal diets can interact to confer complex effects on offspring phenotype. These effects are not simply caused by additive contributions of each parent’s diet. Buescher et al. (2013) similarly demonstrated that maternal and offspring diets interact in ways that are not always additive in *D. melanogaster*, and shape F1 and F2 physiological measures of sugar and fat content, as well as gene regulation linked to lipid metabolism. While intriguing, the broader evolutionary consequences and generality of these maternal-by-paternal diet interactions, and maternal-by-offspring, interactions revealed in each study remain unanswered, since each measured only early-life physiological parameters of offspring, and it is possible that offspring are able to compensate for these early life effects throughout the life course.

Motivated by the questions of whether parental diet effects on offspring fitness phenotypes interact, and whether they may be adaptive for offspring, we tested the relative contributions of variation in maternal and paternal diets to offspring life-history traits (adult lifespan and fecundity) as well as body composition traits (triglycerides, body mass) in the fruit fly *D. melanogaster*. We provided experimental flies with one of two diets that varied in the concentration of sucrose (2.5% and 20%) relative to the other ingredients of yeast, agar, and water. The diets were administered using a fully factorial design, in which the two diets were assigned to mothers, fathers, and offspring in all possible combinations, such that female-male and parent-offspring diet combinations were either matched or mismatched. This design enabled us to test the prediction of whether dietary-mediated intergenerational effects are adaptive (with offspring produced by parents of a matching diet having higher fecundity and lifespan), and whether these effects primarily manifest as maternal or paternal effects, or via interactions between male and female parents.

## Results

### Effects of sucrose treatments on the parental (F0) flies

#### Lifespan and fecundity

The effect of dietary sucrose on longevity of the F0 flies was moderated by sex (F_1,38_= 57, *p* <0.001, Figure 1A, Table S1), with female longevity exhibiting high sensitivity to sucrose (∼35% increase in longevity on a lower sucrose diet relative to the higher sucrose diet). Whereas male longevity *decreased* ∼ 3% on a lower sucrose diet relative to the higher sucrose diet). Notably, the longevity of the F0 males and females was in part dependent on the diet of the flies that they mated with during the brief 96 h cohabitation phase early in life (F_2,37_ ***=*** 3.16, *p* <0.05, Figure 1A, Table S1). Specifically, when the diets of the cohabiting focal and tester flies were matched for sucrose content, the flies lived longer.

**Figure 1.**
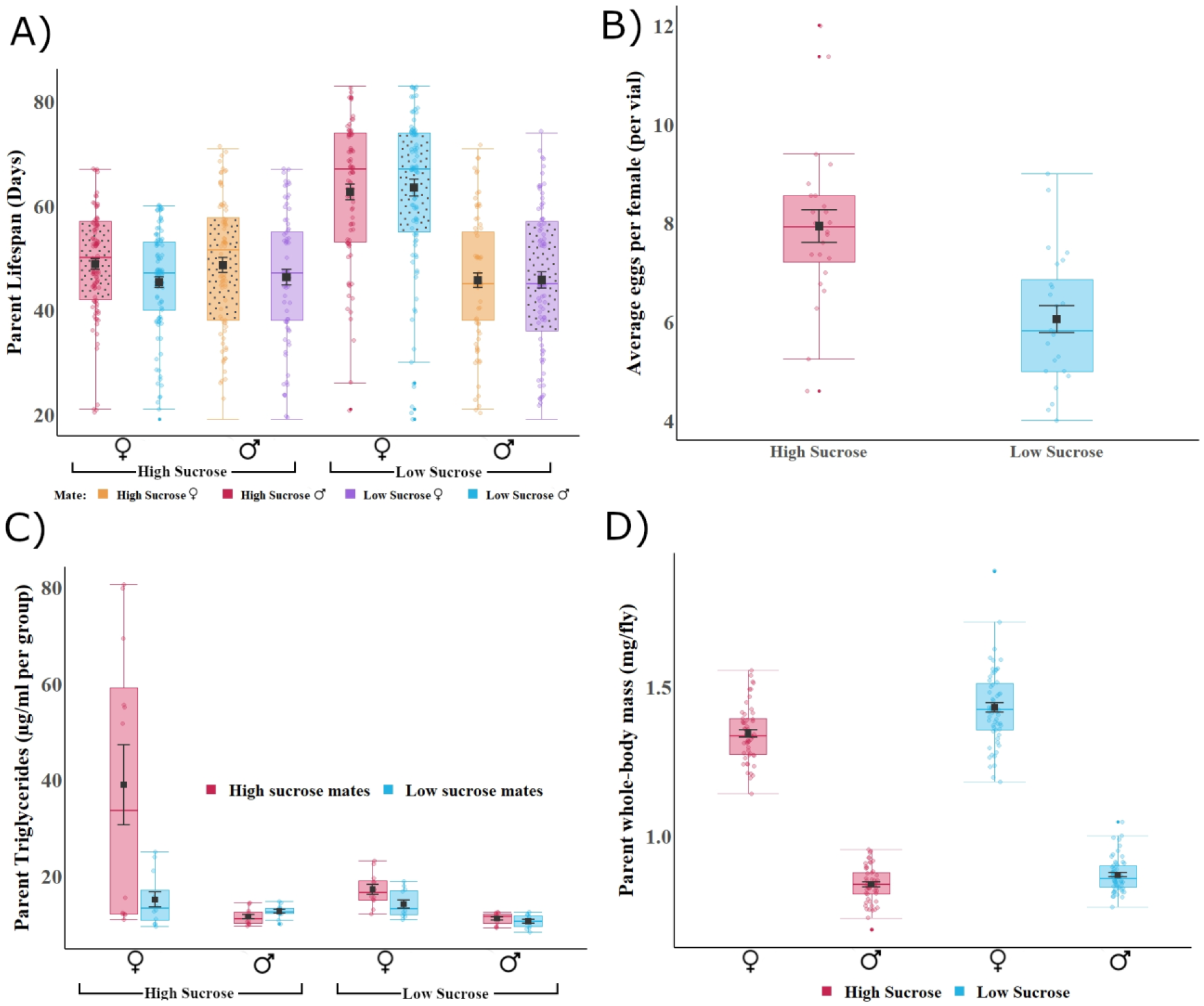
Effects of high sucrose (20% of overall solution), and low sucrose (2.5% of overall solution) on F0 (parent) lifespan, egg production, triglyceride levels, and body mass. Plots show means and standard error bars inside boxplots, showing medians and quartiles. Panel A) Lifespan of F0 flies (y-axis), their diet (x-axis), and diet of the mate (indicated by colour of the boxplots) on lifespan, matching parent diets are indicated by dotted pattern. Panel B) Average eggs per F0 female (y-axis), across both diets (x-axis). Panel C) F0 whole-body triglycerides (y-axis), sex of F0, and their diet (x-axis), with the diet of their mate indicated with colour. Panel D) F0 body mass (y-axis), sex of F0 (x-axis), with their diet indicated with colour.

Females on the lower sucrose diet produced less eggs than those on a higher sucrose diet (F_1,46_ =20.73, *p* <0.001, Figure 1, panel B, and Table S2), but there were no effects of the diet of the males that they mated with on fecundity (F_1,46_ =0.151, *p=*0.699.

#### Lipid and protein measurements

We measured triglyceride levels at two stages in F0 adult flies: prior to mating (at six days post eclosion), and post mating (at nine days post eclosion, at the end of the 96 hour cohabitation period). This enabled us to explore whether the effects of the mate diet on the longevity of flies was potentially underpinned by changes in triglyceride levels following mating.

##### Before mating

Females accumulated more triglycerides than males, both when triglycerides were normalised to protein levels (F_1,12_=7.9 , *p* <0.05, Table S3a), and normalised to body mass (F_1,9_=12.14, *p* <0.01, Table S3b). When normalised to protein content, triglyceride levels were also affected by the interaction between diet and sex of the flies (F_1,12_=11.66 , *p* <0.001). Females fed lower sucrose diets had higher triglyceride levels than those fed higher sucrose (mean ± S.E.: female _low sucrose_ = 3.7µg/ml ± 0.24, female _high sucrose_ = 2.7 µg/ml ±0.19), with the reverse pattern in males (male _low sucrose_ =1.6µg/ml ± 0.24, male _high sucrose_ = 1.22µg/ml ± 0.10).

##### After mating

Triglyceride levels (divided by protein levels) of parents after they mated were affected by their own diet (in a pattern consistent with their pre-mating triglyceride levels) (*lmer* analysis with Kenward-Rogers F test *F*_1,27_***=*** 8.0736, *p* <0.01, Table S4a), and by their sex (*F*_1,27_***=*** 8.3.122, *p* <0.001, Table S4a). Intriguingly, the triglyceride levels after mating were also affected by the diet of their mate (*F*_1,27_***=*** 6.0955, *p* <0.05). These outcomes were unchanged when the data were normalised to body weight, and again triglyceride levels in parents are affected by their sex (F_1,27_= 6.6, *p* <0.05, Table S4b), and by the diet of their mate (F_1,27_= 4.7, *p* <0.05, Table S4b). With the exception of focal males on higher sucrose diets, focal flies of both sexes had higher whole-body triglyceride levels if they mated with a tester fly provided with higher sucrose diet (Figure 1, panel C). This supports the hypothesis that the effect of mate diet on adult longevity may be at least partly mediated by underlying effects on triglyceride levels in the flies.

#### Body mass

Both dietary sucrose content and sex, and their interaction, affected the body mass (measured prior to mating) of the parental F0 flies (F_1, 206_ 5.27, *p* <0.05 Table S5). Flies assigned to the lower sucrose diet were heavier compared to flies assigned to the higher sucrose diet, with the difference in body mass across the two diets greater in females than males (Figure 1, panel D).

#### Feeding behaviour

Sucrose content did not affect feeding behaviour (number of probosis extentions onto the food) for the F0 flies, but an interaction between age and sex affected feeding behaviour (F_1,118_ 12.14, *p* <0.001, Table S6, Figure S1), with females feeding more than males at days 8 and 11, but with sex differences dissipating at later life stages.

### Effects of parental diets on offspring (F1)

#### Lifespan

The lifespan of F1 flies was shorter for flies produced by parents whose diets were matched for sucrose content than those born to parents whose diets were mismatched (*lmer* analysis, *F*_*1,103*_ ***=*** 4.82, *p* <0.05, Table S7, supplementary material, Figure 2, panel A). Dietary sucrose intake of the F1 females also directly affected their longevity in a manner that mimicked the effects in the F0 flies; high sucrose diets greatly decreased the lifespan of females relative to the low sucrose diet. Intriguingly, the pattern was reversed in F1 males, with males on the higher sucrose diet outliving those on the lower sucrose diet (*F*_*1,103*_ ***=*** 249.37, *p* <0.001, Table S7, supplementary material). There was no interaction between offspring diet and either the maternal or paternal diet, indicating no signatures of an adaptive intergenerational effect for longevity.

**Figure 2.**
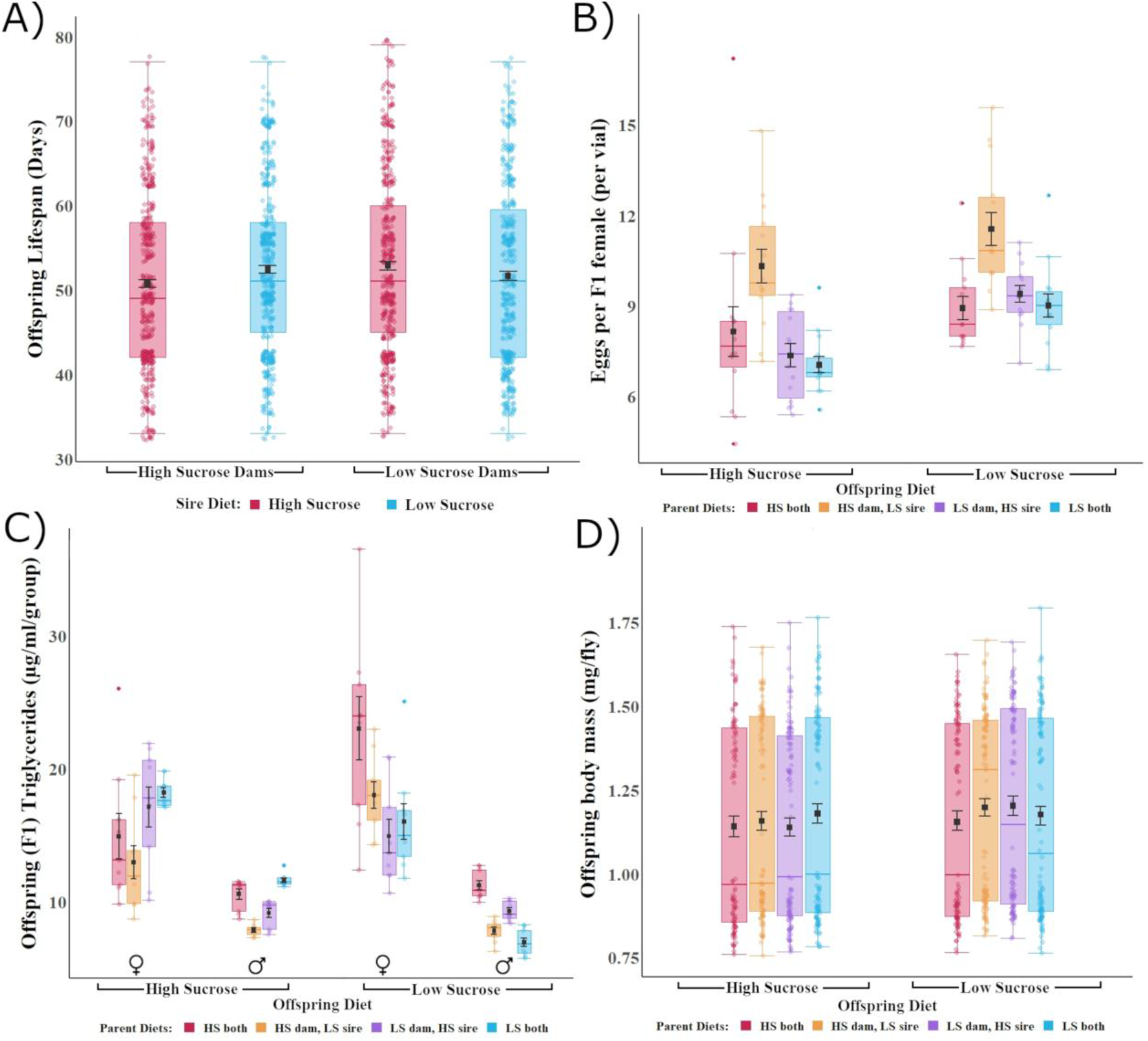
Effects of high sucrose (20% of overall solution), and low sucrose (2.5% of overall solution) on F1 (offspring) lifespan, egg production, triglyceride levels, and body mass. Plots show means and standard error bars inside boxplots, showing medians and quartiles. Panel A) Lifespan of F1 flies (y-axis), their dam’s diet (x-axis), and diet of their sire indicated by colour of the boxplots. Panel B) Average eggs per F1 female (y-axis), their diets (x-axis), and the parental diet combination indicated by colour. Panel C) F1 whole-body triglycerides (y-axis), sex of F1, and their diet (x-axis), with the diet combination of their parental indicated with colour. Panel D) F1 body mass (y-axis), diet of F1 (x-axis), with the parental diet combination indicated with colour.

#### Fecundity

On average, female offspring flies assigned to a low sucrose diet generally oviposited more eggs than those on a high sucrose diet (Figure 2, panel B). This general pattern differed to that observed for fecundity of the F0 females, where flies on the higher sucrose had a greater egg output. Notably, F1 egg output was shaped by a complex interaction between the F1 diet, maternal diet, and paternal diet (*F*_*2,103*_ =8.02, *p* <0.001, Table S8, supplementary material), albeit the pattern was not in the direction predicted under a scenario of an adaptive intergeneration effect, in which matched parental-offspring combinations would be expected to outperform mismatched combinations. Rather, we observed a large effect of one particular parental diet combination on intergenerational fecundity, in which F1 fecundity was higher when the dams were exposed to high sucrose, but the sires low sucrose (Figure 2B).

#### Whole-body triglycerides

A complex interaction between F1 diet, sex, and the diets of both parents shaped whole-body triglyceride levels in the F1 offspring (*lmer* analysis with Kenward-Roger’s method *F*_*3,32*_***=*** 3.93, *p* <0.01, Fig 2C, Table S9). Triglyceride levels were lower when the F1 female diet was matched to the maternal diet, but these patterns were not observed for F1 males (Figure 2, panel C). Both F1 females and males assigned to the low sucrose treatment were generally characterised by low triglyceride content when produced by parents who had both consumed low sucrose, and high triglyceride content when produced by parents that had both consumed high sucrose. Conversely, F1 females and males that had been provided with a high sucrose diet were characterised by the highest triglyceride content when produced by parents that had both consumed low sugar.

#### Body mass

An interaction between the three diets, that is, the sucrose content of the dam, sire, and offspring affected the body mass of the F1 offspring (F_2,86_ 4.50, *p* <0.05, Table S10, supplementary material). Flies that consumed the lower sucrose diet, but who were produced by parents whose diets were matched to each other (i.e. either both parents consumed high sugar, or both consumed low sugar) weighed less than flies whose parents consumed diets that were mismatched for sucrose (Figure 2, panel D).

#### Feeding behaviour

Sucrose content of either the parental diets or the offspring diets did not have a significant effect on offspring feeding behaviour (number of probosis extentions onto the food). However, feeding behaviour differed across the sexes, with females feeding more than males, especially during peak reproductive periods of eight and eleven days post eclosion (F_1,43_ 24.85, *p* <0.001, Table S11, Figure S1, supplementary material).

#### Correlations

In our dataset, we tested associations between triglyceride levels, fecundity, and lifespan, and found them non-significant in all cases, indicating that if associations exist they are weak. Further information can be found in the supplementary material.

## Discussion

We varied the concentration of sucrose relative to all other nutrients in the diet of female and male *D. melanogaster*, across two generations, and examined the response in the expression of both life-history and physiological traits. We used a fully factorial design in which sires, dams, and their offspring were provided with diets that were either higher (20% of overall solution) or lower (2.5% of overall solution) in relative sucrose, such that combinations of parental diets and parent-offspring diets were either matched or mismatched, in all possible combinations. This design provided an opportunity to screen for adaptive dietary-mediated anticipatory parental effects on offspring phenotypes, and an opportunity to explore relative maternal and paternal contributions to offspring performance following dietary manipulation.

Our study revealed several new findings. First, although we detected parent-by-offspring diet interactions on offspring fecundity, triglyceride content, and body mass, rarely were these in a direction consistent with predictions of the effects being anticipatory and adaptive. Moreover, generally these interactions were complex, with offspring trait values contingent on the diets of all interacting parties – sire, dam, and offspring. As such, dietary-mediated parental contributions to offspring phenotypes were typically non-additive rather than cumulative, with particular combinations of mismatched dam-sire diets conferring heightened trait expression in offspring. Second, we identified unexpected dietary-mediated effects on the lifespan of both F0 and F1 generations. Notably, the lifespan of F0 flies was affected by the diets of their mates, with flies paired to mates that had been fed a diet matched in sucrose content to their own diets exhibiting longer lifespan than flies paired to mates fed a mismatched diet. Remarkably, however, while interacting dams and sires whose diets were sucrose-matched enjoyed longer lives, their offspring suffered a longevity disadvantage relative to offspring produced by dams and sires whose diets were mismatched for sucrose. This suggests potential for a parent-offspring conflict over optimal dietary sucrose ingestion.

### Evidence for anticipatory parental effects

The key prediction underpinning the hypothesis of anticipatory parental effects is that components of offspring fitness will be higher when the offspring environment matches the parental environment—a prediction that has been most often tested in the context of maternal effects on offspring fitness (Mousseau & Fox, 1998; Uller et al., 2013; Yin et al., 2019). Testing this requires a particular experimental design whereby parents and offspring are exposed to matched and mismatched environments, and predicts that matched combinations (between parent and offspring) will result in the expression of offspring trait values that maximise fitness, and are hence adaptive, relative to mismatched combinations (Burgess & Marshall, 2014; Uller et al., 2013). While this prediction has received support from both classic and recent studies that used match-mismatch designs (Agrawal, Laforsch, & Tollrian, 1999), meta-analyses aimed at synthesizing patterns across species have however produced only weak evidence that such effects exist generally (Radersma et al., 2018; Sánchez-Tójar et al., 2020; Uller et al., 2013; Yin et al., 2019). Some evidence suggests the failure to detect general effects might be due to methodological deficiencies across studies (Burgess & Marshall, 2014; Uller et al., 2013), for example, a failure to test intergenerational outcomes in both matched and mismatched combinations, or the inability to partition anticipatory effects from condition-dependent parental effects. More generally, partitioning condition-dependent parental effects from cases that are genuinely anticipatory is likely to represent an ongoing challenge. Even in cases where researchers employ match-mismatch designs, given that condition-dependent effects may be context dependent in some cases and mimic patterns expected under the prediction of an anticipatory scenario. For example, this might occur in the case of “silver-spoon” scenario, in which parents in better condition may produce offspring in better condition, compared to their lower conditioned counterparts, but only under certain environmental conditions (Bonduriansky & Crean, 2018; Engqvist & Reinhold, 2016).

Currently, it is unclear how often anticipatory parental effects might be triggered by environmental heterogeneity in the quality of food available to individuals. Many studies investigating dietary-mediated intergenerational or transgenerational effects in model organisms (in the context of nutritional and metabolic programming) have not implemented the requisite full factorial designs required to test the hypothesis (Buescher et al., 2013; Huypens et al., 2016; Matzkin, Johnson, Paight, & Markow, 2013; Oldham, 2011; Öst et al., 2014; Polak et al., 2017). Moreover, meta-analyses conducted to date have not sought to disentangle the relative strength of different classes of environment (e.g. dietary, climatic, pathogenic) on the magnitude of effect sizes associated with inter- or transgenerational anticipatory effects (Sánchez-Tójar et al., 2020; Uller et al., 2013; Yin et al., 2019). Notwithstanding, heterogeneity in the food environment would seem to be an excellent candidate to drive intergenerational anticipatory effects, given that macronutrient availability is likely to be relatively stable across generations for many species, and such predictability is a key theoretical requirement underpinning the evolution of anticipatory parental effects (Burgess & Marshall, 2014). We thus tested for anticipatory parental effects in the context of dietary sucrose environments, testing whether offspring lifespan, fecundity, and physiology were sensitive to parent-offspring interactions.

While we found that fecundity, triglyceride content, and body mass are sensitive to such interactions, only the patterns for female triglyceride content were consistent, and only weakly so, with the prediction that parent-offspring matches might result in a superior phenotype. In particular, female offspring who had been assigned to higher sucrose diets had lower triglyceride contents if their mothers had also been assigned to the higher sucrose treatment. Similarly, female offspring on a lower sucrose diet had a lower triglyceride content if their mothers were also provided lower sucrose. These signatures of intergenerational anticipatory effects were not evident in male offspring, and were only transmitted through dams. Whether or not these signatures of anticipatory effects are adaptive would depend generally on the association between triglyceride levels and fitness. The association would need to assume that low triglyceride levels confer higher lifetime reproductive success, and there is some evidence in *D. melanogaster* to suggest this may be the case. Studies that use *Drosophila* as a model for studying effects of diet, obesity, and exercise, have shown that heightened activity, simulating exercise, in flies leads to reductions in triglyceride content (Sujkowski & Wessells, 2018), and a previous study has shown a sharp decline in female fecundity with increasing triglyceride levels (Skorupa, Dervisefendic, Zwiener, & Pletcher, 2008). Notwithstanding, in our dataset associations between triglyceride levels, fecundity, and lifespan were non-significant in all cases, indicating that if associations exist they are weak. Thus, we conclude that if mechanisms of anticipatory parental effects, regulated by dietary sucrose variation, are at play in this species, the effects are weak and dwarfed by complex and non-additive maternal by paternal diet interactions, we discuss in more detail below.

### Non-additive maternal and paternal effects on both parent and offspring lifespans

We observed a contradictory pattern across generations, whereby dietary matching between males and females extended lifespan of the interacting flies, but reduced lifespan among their offspring. Such a result is intriguing and indicative of potential antagonism between generations in terms of the optimal macronutrient balance underpinning expression of key fitness-related traits. Our result is concordant with results of two recent transgenerational dietary restriction studies in *C. elegans*. In these studies, the authors revealed what they termed ‘missing costs’ of dietary restriction, demonstrating that a parental optimum for temporary fasting (restricted food for 6 days) that increased their own survival, reproduction, and heat tolerance, incurred negative effects on offspring fitness, and notably increased the mortality risk in the great-grandparental (F3) generation. Female offspring produced by long-lived mothers that had fasted, had lower lifetime reproductive output, smaller body size, and slower development than daughters from mothers that had not fasted (Mautz, Lind, & Maklakov, 2020; Ivimey-Cook et al., 2020.). Our results reaffirm the contention that intergenerational effects, mediated by dietary restriction or modification to macronutrient balance, may differ not only in their relative magnitude from one generation to the next, but also in their direction. These results highlight a research avenue that deserves further attention.

The observation that an individual’s lifespan is shaped in part by the diet of their mate is remarkable, especially given that the flies used in our experiments only cohabited for a period of four days early in life. This begs the question of what underlying physiological processes may mediate these effects. One possibility is that the diets of flies directly affected their condition, and subsequently shaped levels and intensity of sexual interaction between males and females. Increases in sexual interaction have been shown to decrease lifespan of female *D. melanogaster* (Bretman & Fricke, 2019; Liddle, Chapman, Partridge, Kalb, & Wolfner, 1995; Wigby & Chapman, 2005) and these effects also carry over to the next generation, resulting in a decreased lifespan among offspring (Dowling, Williams, & Garcia-Gonzalez, 2014). A previous study has also confirmed that dietary quality of males (levels of yeast in the diet) affects their reproductive competitiveness under sexual selection, in a non-linear pattern (Fricke, Bretman, & Chapman, 2008). Moreover, the effects we observed may be partly mediated by triglyceride levels of the interacting flies. Our analyses of triglyceride levels of flies provide some insight, since females who cohabited and mated with males subjected to higher sucrose diets had higher whole-body triglyceride levels post-mating when compared to those who cohabited with males provided with lower sucrose mates. This suggests some capacity for males to directly alter the physiological status of their mates through transfer of seminal proteins during mating (Chapman, Liddle, Kalb, Wolfner & Partridge, 1995). The capacity for dietary variation to mediate patterns that shape the outcomes of interacting phenotypes—phenotypes that are partly mediated by non-genetic effects among conspecifics— and the role of triglycerides in moderating effects on female life history following mating, warrants further investigation.

Finally, we note that the nature of the parent-offspring diet interactions we observed were generally complex and contingent on the diets of interacting flies. For example, the main determinant of female offspring fecundity was an interaction between maternal and paternal diet, which affected female offspring fecundity independently of the diet of the female offspring. In particular, female F1 offspring produced by mothers fed higher sucrose and fathers fed lower sucrose had substantially higher fecundity than female offspring produced by any other combination of parental diet. These interactions suggest that effects of maternal and paternal diet on offspring phenotypes will not be simply cumulative, but rather the result of non-additive interactions.

### Future directions

We suggest that future work expand the range of diet treatments that we used here to investigate whether the antagonistic effects mediated by dietary matching that we observed across generations are specific to the dietary treatments we used, or whether they can be generalised across a broader range of protein to carbohydrate ratios and caloric contents. There have been recent calls for such experiments that utilise the nutritional geometric framework within a transgenerational context (Bonduriansky & Crean, 2018). Additionally, our study measured reproductive consequences of the different dietary treatments in females only, and over a short period early in adult life. We suggest that future studies focus on the intergenerational effects of diet on reproductive success. This is important because negative intergenerational effects that we reported on F1 lifespan may indeed be adaptive if accompanied by overall increases in reproductive output across the F1 lifespan. Exploration of effects beyond the F1 would facilitate interpretation of whether the patterns reported here are more likely mediated by direct condition-transfer from parents to offspring, or via epigenetic mechanisms that are more likely to be anticipatory in nature (Sánchez-Tójar et al., 2020). Finally, we note that the insights gained here from the study of fruit flies may have relevance to mechanisms underpinning nutritional programming in mammalian systems including humans, given that many of the genes and metabolic pathways involved in nutrition, obesity, and aging are generally conserved (Camilleri-Carter et al., 2019).

## Methods

### Study species and generating experimental flies

We sourced flies from a large laboratory population of *D. melanogaster* (Dahomey), originally sourced from Benin, West Africa (Puijk, 1972). The flies have been maintained in large population cages, with overlapping generations in the Piper laboratory at Monash University since 2017, and prior to that in the Partridge laboratory at University College London (Mair, Piper, & Partridge, 2005). Prior to the beginning of the experiment, we collected ∼3000 eggs from the cages, and distributed them into 250mL bottles containing 70mL of food, at densities of 300-320 adults per bottle. Food comprised 5% sucrose (50 grams sucrose, 100 grams yeast, 10 grams agar per 1 litre solution with an estimated protein to carbohydrate [P:C] ratio of 1:1.9, and 480.9kcal per litre, (see Table S12, and Figure S2, supplementary material for further diet details). Each generation, we admixed adult flies, emerging from across different bottles, together before redistributing back into bottles at a density of 300-320 adults per bottle, repeating this for 7 generations. To control for potential sources of variation in their environment during these 7 generations, we controlled the age of flies at the time of ovipositioning (all flies were within 24 h of eclosion into adulthood when producing the eggs that propagated the subsequent generation) and the egg density within each bottle (∼300 eggs per bottle).

### Dietary treatments

The diet media we used consists of sucrose, autolysed brewer’s yeast powder (sourced from MP Biomedicals SKU 02903312-CF), and agar (grade J3 from Gelita Australia), as well as preservatives—propionic acid, and nipagin. We prepared two dietary treatments, differing in relative sucrose concentration; 2.5% sucrose (that we refer to as a lower sucrose treatment, relative to the 5% concentration usually provided to the population of flies used in this experiment), and 20% sucrose (that we refer to as a higher sucrose treatment) of overall food solution. The 2.5% sucrose diet contains 25 grams of sucrose, 100 grams of yeast and 10 grams of agar per litre of food prepared, with an estimated P:C ratio of 1:1.4 and 380.9kcal per litre of food. The 20% sucrose treatment contains 200 grams of sucrose, 100 grams of yeast, and 10 grams of agar per litre of food prepared, with an estimated P:C ratio of 1:5.3 and 1080.9kcal per litre of food. The diets thus differed not only in sucrose concentration, but overall macronutrient balance and their total caloric content. The high sucrose concentration was selected based on preliminary experiments that we conducted, which caused the flies to accumulate a higher content of triglycerides but still allowed for offspring production and viability (relative to other sucrose levels), consistent with results from previous work in *D. melanogaster* (Buescher et al., 2013; Skorupa et al., 2008). All diets contained 3ml/l of propionic acid and 30ml/l of a Nipagin solution (100g/l methyl 4-hydroxybenzoate in 95% ethanol) and were prepared according to the protocol described in (Bass et al., 2007). Each vial is 40mL and contained 7mL of food.

### Experimental design

Male and female virgin flies were assigned to one of the two dietary treatments prior to mating (we refer to this generation of flies as F0), and then the offspring produced (we refer to this as the F1 generation) were also assigned to one of the two treatments. All possible combinations of dam × sire × offspring diet treatment were represented (= 2 × 2 × 2 = 8 combinations). Specifically, we collected flies of the F0 generation as virgins and placed them in vials of 10 flies across 30 vial replicates per treatment (1300 flies, 650 of each sex), in their respective sexes, onto either the high sucrose (20%) or the low sucrose (2.5%) diets for the first 6 days of their adult life. We transferred flies to vials containing fresh food of the designated diet every 48 hours during this 6 day period.

At day 6, we randomly sampled six vials from each treatment, and snap froze (using liquid nitrogen) the flies of these vials, storing them at -80°C for later triglyceride and body-weight assays. Cohorts of flies in the remaining vials then entered a cohabitation phase to enable female and male flies to mate. Cohorts of males and female flies were combined, in vials of 10 pairs, in each of all four possible diet combinations: Lower sucrose females × lower sucrose males; higher sucrose females × higher sucrose males; lower sucrose females × higher sucrose males; higher sucrose females × lower sucrose males. During this phase, flies cohabited for 96 hours, allowing them to mate. They were transferred to a new vial with fresh food of standard 5% sucrose diet every 24 hours during this time.

Following the cohabitation phase of 96 h, the F0 flies were separated back into their respective sex-specific cohorts, and placed back onto the high or low sucrose diets that they were originally assigned prior to the cohabitation phase, in vials of 20 flies. Flies of these vials were then monitored for longevity (the longevity assay is described below). The vials from the 6 day old F0 flies (i.e., the vials from day 1 of the 96 h cohabitation phase) were retained, and the eggs that had been laid by females of the respective vials were trimmed to 80 per vial (by removing excess eggs with a spatula). The remaining eggs were left to develop into adult offspring over 10 days at 25°C (on a 12:12 light/dark cycle in a temperature-controlled cabinet; Panasonic MLR-352H-PE incubator). These adult flies constituted the F1 offspring in the experiment, all F1 were reared on standard media (5% sucrose). We collected virgin F1 adults from each of the four combinations of parental diet treatments, and placed them in their respective sexes in vials of 10 flies, across 30 vial replicates per treatment per sex (2400 flies). We then assigned these F1 flies, produced by each dietary treatment combination of F0 flies, to either the lower sucrose or higher sucrose diet. At day 6 of adulthood, we snap froze F1 flies of six randomly chosen vials per dam × sire × offspring diet combination. On the same day, 10 virgin focal F1 flies were placed together with 10 *tester* flies of the opposite sex (age standardised), collected from the Dahomey stock population, entering into a cohabitation phase of 96 h (during which time the number of eggs laid by females of each vial was assessed). After 96 hours flies were separated again into their respective sexes (in vials of 20 flies), and assigned back onto either the lower sucrose or higher sucrose diets that they had been on prior to cohabitation, and a longevity assay carried out.

### Longevity

We scored the longevity of experimental flies of both parental (F0) and offspring (F1) generations. Cohorts of each sex were assayed separately. Each vial in the assay commenced with 20 flies each, and we included six vial replicates per treatment combination (dam × sire) for the F0 (700 flies), and seven vial replicates per treatment combination (dam × sire × offspring) for the F1 (2000 flies). The number of dead flies per vial was scored three times per week (Monday, Wednesday, Friday), and surviving flies at each check transferred to vials with fresh food of the assigned diet treatment—until all flies were deceased. During the lifespan assay, vials were stored in boxes of up to 100 vials each, which were moved to randomised locations in the (25°C) control temperature cabinet every few days to decrease the potential for confounding effects of extraneous and random environmental variation from affecting the results.

### Fecundity

We measured the egg output of female flies from generations F0 and F1 at eight days following the females’ eclosion to adulthood, and used these egg counts as a proxy of female fecundity. On day eight, female flies oviposited for a 23 hour period, and were then transferred to fresh vials. Day eight was selected because fecundity over 24 hours at this age has been shown to correlate with total lifetime fecundity of females in this Dahomey population (Nguyen & Moehring, 2015). Additionally, day eight aligns with the peak period in reproductive output in the species (Bass et al., 2007) and early, short term measures of reproduction of between one and seven days can be used to accurately predict total lifelong fecundity in *D. melanogaster* (Nguyen & Moehring, 2015). Moreover, prior data shows that modification of sucrose concentrations does not alter the timing of the reproductive peaks between treatments [(Bass et al., 2007) for more information see Figure 8 in the supplementary information]. For the F0 generation, we counted eggs from 12 vial replicates per sire x dam combination, each containing 10 female flies, that had been mated with 10 male flies, across 2 different sucrose levels (2.5% and 20% sucrose), as above. We also counted eggs from F1 female flies, sampling 14 vial replicates per sire × dam × offspring combination, each containing 10 focal females (females from the experiment) combined with 10 tester male flies.

### Feeding behaviour

A separate experiment was set up to assess the feeding behaviour of the adult flies. Flies from the same wild type Dahomey population were used as in the previous fecundity and longevity assays, and kept under the same conditions as they were for previous assays (in the same parental and offspring diet combinations as described above), with the exception of the number of flies per vial. For this assay, flies were kept in vials of five individuals, separately by sex, except during the 96-hour cohabitation window when they were kept in vials of 10 flies, five males and five females. These flies were transferred to new food every 24 hours, always transferring them to new food the night before an observation.

We measured feeding behaviour of the flies of the F0 and F1 generations of each combination of diet treatment, using previously reported protocols of Wong, Piper, Wertheim, and Partridge (2009). Feeding behaviour of the flies of each vial was observed over a two-hour observation period, commencing at 10 am, including 30 minutes of time acclimating flies to the observer’s presence. Observations were run four times for each vial over the first three weeks of adult life at 8, 11, 15 & 17 days of age for the parental generation, and 8, 11, 15 & 20 days of age for the offspring generation. This was performed for each of the two different dietary treatment levels, (2.5% and 20% sucrose), and for each of the parental dietary treatment combinations (dam × sire × offspring), to address whether the flies moderated their feeding behaviour according to the dietary treatment they were subjected to, or the treatment of their parents.

### Body weight

Adult flies that had been snap frozen at six days post eclosion were individually weighed with a Mettler Toledo ultra-microbalance (Model: XP2U/Z). In the F0 generation, 220 (110 female, 110 male) flies were weighed from the two different dietary treatments. In the F1 generation, 879 flies (439 males, 440 females) were weighed from the two dietary treatments, (219-220 per parental diet treatment combination).

### Lipids and protein

Whole-body triglyceride levels were measured in adult flies from the (F0) parental generation (six days of adult age and corresponding to six days on the dietary treatment, prior to mating) and normalized to body weight (full protocols reported in the Supplementary Material). A separate experiment was set up to generate additional samples of the (F0) parental generation, this time, after they had cohabited for 96 hours, and snap frozen at 9 days following eclosion. Flies were kept under the same conditions as they were for previous assays (as described above). We also added an additional assay to measure whole-body protein. This was done to determine whether the protein and triglyceride content of the (F0) parental flies would be altered after mating, using protein as a proxy for the amount of metabolically active tissue.

Six biological replicates (from different vials) per treatment level were used to conduct the triglyceride and protein assays in the F0, as well as three technical replicates (repeated aliquots from the same sample of adult flies. For the F1 generation, three biological replicates per treatment level, with three technical replicates per biological replicate were used. Five female flies and eight male flies respectively, were used for each biological replicate in the assay.

### Statistical Analyses

We used R (Version 3.6.1) and RStudio (Version 1.2.1335) (R Core Team, 2019) for statistical analyses. We modelled dietary effects on both the F0 and F1 flies, running separate models for each generation, and each trait (lifespan, fecundity, feeding behaviour, body weight, and triglyceride level). We fitted linear mixed effects models, using the R package lme4 (Bates, Mächler, Bolker, & Walker, 2015), to test the effects of fixed factors parental diet, offspring diet, mate diet, and sex, and possible interactions between these fixed effects, on lifespan of the F0 flies (offspring diet and offspring sex was not included in F0 analyses) and F1 flies respectively. We included the vial identification number as a random intercept in the longevity models. Since we monitored flies for lifespan thrice weekly (Monday, Wednesday, and Friday), the age of recorded death for each individual fly was estimated within a margin of 72 hours (for example, a lifespan of 30 days indicates that a fly died between 27-30 days post eclosion).

To test the effects of parental diet, offspring diet, mate diet and sex on female fecundity, we fit a linear model to the egg output data for both generations. We included parental diet, offspring diet, mate diet and sex as fixed effects, and we explored interactions between these factors. The fecundity models only included one observation per vial because we counted the overall number of eggs laid per group of females of a given vial, and divided by the number of females in the vial, to derive an average per female, and therefore no random intercept was required in these models.

We first fit full models to the data, including all fixed effects and their interactions (up to the level of second order interactions). We reduced each model down to a final (minimum adequate) model using an approach based on parsimony reduction, in which the least-significant terms were removed sequentially, starting with the highest order interactions. We tested whether the reduction of each term led to a significant change in the deviance between models with log-likelihood ratio tests, and an alpha criterion of 0.05. The final models for lifespan, body weight, feeding behaviour, and triglyceride level were fit by restricted maximum likelihood, applying type III F tests with Kenward-Roger’s approximation of degrees of freedom. Fecundity measures were fit using F tests and Type III sums-of-squares ANOVA. We visually inspected diagnostic plots for the linear mixed effect models, to ensure that the assumptions of normality and equal variances were met. To investigate the relationships between fitness indicating traits (egg production and lifespan) with body composition (body mass and triglyceride content) we calculated Pearson’s pairwise correlations and then bias corrected bootstrapped confidence intervals.

## Supporting information

All supplemental information

## Author contributions

TLCC, DKD, MDWP & RLR conceptualised the study and designed the experiment, TLCC planned and carried out the experiment, wrote the first draft of the manuscript, and TLCC, DKD, MDWP & RLR contributed to the writing and editing of the subsequent drafts.

## Acknowledgements

We are grateful for the help we received from laboratory assistants, Pavani Manchanayake, James Wang, Skye Bulka, and Natalie Wagan. Additional invaluable assistance was provided by Caleb Carter, and Amy Dedman,

