## Supplementary material for "Maternal and paternal sugar consumption interact to modify offspring life history and physiology": All supplemental information

#### Results

#### F0 Lifespan

**Table S1.** Table S1. Effects of diet of the focal fly (Diet), sex, and diet of the mate (Mate) on longevity of the focal flies. General Linear Mixed Model, with fixed effects parameters calculated via F test with Kenward-Rogers approximation of degrees of freedom, and variance associated with random effect (Vial identity) estimated via REML.

*** indicates significance at $p < 0.001$, ** indicates significance at $p < 0.01$, * indicates significance at $p < 0.05$.

| ***Fixed Effects*** | ***F*** | ***p-value*** |
|---|---|---|
| (Intercept) | 1031.32 | < 0.001 *** |
| Diet | 92.88 | < 0.001 *** |
| Sex | 1.73 | 0.185 |
| Mate | 2.07 [a] | 0.122 |
| Diet: sex | 57.47 | < 0.001 *** |
| Diet : mate | 3.16[a] | < 0.05 * |
| ***Random Effects*** | ***Variance*** | |
| Vial identification | 0.0 | |
| Residual | 200.8 | |

[a]df=2, all other df=1, df res=38

#### F0 Fecundity

**Table S2.** Statistical results (analysis of deviance) of the linear model (Gaussian error distribution) after model reduction for predictors of female (F0) egg output.
*** indicates significance at $p < 0.001$, ** indicates significance at $p < 0.01$, * indicates significance at $p < 0.05$.

| ***Fixed Effects*** | ***Sum Sq*** | ***F*** | ***p-value*** |
|---|---|---|---|
| (Intercept) | 1507.02 | 758.84 | < 0.001 *** |
| Female diet | 41.94 | 20.73 | < 0.001 *** |
| Male diet | 0.31 | 0.151 | 0.699 |

all *df* =1, df res=46

#### F0 TAG & Protein

**Table S3a. A**nalysis of deviance table with Kenward-Rogers F test, linear mixed model (Gaussian error distribution) after model reduction for predictors F0 whole-body triglycerides, normalised by protein—before mating.
*** indicates significance at $p < 0.001$, ** indicates significance at $p < 0.01$, * indicates significance at $p < 0.05$.

| ***Fixed Effects*** | ***F*** | ***p-value*** |
|---|---|---|
| Intercept | 115.28 | < 0.001 *** |
| Diet | 0.48 | 0.50 |
| Sex | 7.87 | < 0.05 * |
| Diet : Sex | 11.65 | < 0.01 ** |
| ***Random Effects*** | ***Variance*** | |
| Vial identification | 0.1475 | |
| Residual | 0.01126 | |

*All df* =1, df res=12

**Table S3b. A**nalysis of deviance table with Kenward-Rogers F test, linear mixed model (Gaussian error distribution) after model reduction for predictors F0 whole-body triglycerides, normalised by body weight—before mating.
*** indicates significance at $p < 0.001$, ** indicates significance at $p < 0.01$, * indicates significance at $p < 0.05$.

| ***Fixed Effects*** | ***F*** | ***p-value*** |
|---|---|---|
| Intercept | 28.09 | < 0.001 *** |
| Diet | 0.88 | 0.37 |
| Sex | 12.14 | < 0.01 ** |
| ***Random Effects*** | ***Variance*** | |

| | |
|---|---|
| Vial identification | 0.000 |
| Residual | 0.000 |

*All df* =1, df res=9

**Table S4a.** Analysis of deviance table with Kenward-Rogers F test, linear mixed model (Gaussian error distribution) after model reduction for predictors F0 whole-body triglycerides, normalised by whole-body protein—after mating.

*** indicates significance at $p < 0.001$, ** indicates significance at $p < 0.01$, * indicates significance at $p < 0.05$.

| ***Fixed Effects*** | ***F*** | ***p-value*** |
|---|---|---|
| Diet | 8.0736 | < 0.01 ** |
| Sex | 8.3122 | < 0.001 *** |
| Mate diet | 6.0955 | < 0.05 * |
| ***Random Effects*** | ***Variance*** | |
| Vial identification | 0.00026 | |
| Residual | 0.00027 | |

*All df* =1, df res=27

**Table S4b.** Analysis of deviance table with Kenward-Rogers F test, linear mixed model (Gaussian error distribution) after model reduction for predictors F0 whole-body triglycerides, normalised by body weight—after mating.

*** indicates significance at $p < 0.001$, ** indicates significance at $p < 0.01$, * indicates significance at $p < 0.05$.

| ***Fixed Effects*** | ***F*** | ***p-value*** |
|---|---|---|
| Diet | 8.0736 | < 0.10 |
| Sex | 8.3122 | < 0.05 * |
| Mate diet | 6.0955 | < 0.05 * |
| ***Random Effects*** | ***Variance*** | |
| Vial identification | 0.0000017 | |
| Residual | 0.00000033 | |

*All df* =1, df res=27

#### F0 Body mass

**Table S5.** Statistical results (analysis of deviance) of the linear mixed model (Gaussian error distribution) after model reduction for predictors of whole-body weight for the parental generation. *** indicates significance at $p < 0.001$, ** indicates significance at $p < 0.01$, * indicates significance at $p < 0.05$.

| *Fixed Effects* | *F* | *p-value* |
|---|---|---|
| (Intercept) | 7128.55 | < 0.001 *** |
| Diet | 15.51 | < 0.001 *** |
| Sex | 862.63 | < 0.001 *** |
| Diet: sex | 5.28 | < 0.05 * |
| ***Random Effects*** | ***Variance*** | |
| Vial identification | 0.000 | |
| Residual | 0.007 | |

*All df* =1, df res=206

#### F0 Feeding behaviour

**Table S6.** Statistical results (analysis of deviance with Kenward-Rogers's method) of the linear mixed model (Gaussian error distribution) after model reduction for predictors of feeding behaviour for the parental generation. *** indicates significance at $p < 0.001$, ** indicates significance at $p < 0.01$, * indicates significance at $p < 0.05$.

| *Fixed Effects* | *F* | *p-value* |
|---|---|---|
| Diet | 1.36 | 0.2506 |
| Sex | 19.88 | < 0.001 *** |
| Age | 18.32 | < 0.001 *** |
| Sex : Age | 12.14 | < 0.001 *** |
| ***Random Effects*** | ***Variance*** | |
| Vial identification | 0.8546 | |
| Residual | 5.5823 | |

*All df* =1, df res=118

#### F1 Lifespan

**Table S7. A**nalysis of deviance table with Kenwood-Rogers F test, linear mixed model (Gaussian error distribution) after model reduction for predictors of offspring age.
*** indicates significance at $p < 0.001$, ** indicates significance at $p < 0.01$, * indicates significance at $p < 0.05$.

| *Fixed Effects* | *F* | *p-value* |
|---|---|---|
| (Intercept) | 2926.96 | < 0.001 *** |
| Offspring diet | 381.54 | < 0.001 *** |
| Offspring sex | 14.99 | < 0.001 *** |

| | | |
|---|---|---|
| Maternal diet | 1.92 | 0.154 |
| Paternal diet | 1.47 | 0.209 |
| Offspring diet : offspring sex | 249.37 | < 0.001 *** |
| Maternal diet : paternal diet | 4.82 | < 0.05 * |
| ***Random Effects*** | ***Variance*** | |
| Vial identification | 3.2 | |
| Residual | 122.1 | |

All *df =1*, df res=103

### F1 Fecundity

**Table S8.** Statistical results (analysis of deviance) of the linear model (Gaussian error distribution) after model reduction for predictors of female offspring egg output.
*** indicates significance at *p* < 0.001, ** indicates significance at *p* < 0.01, * indicates significance at *p* < 0.05.

| ***Fixed Effects*** | ***Sum Sq*** | ***F*** | ***p-value*** |
|---|---|---|---|
| (Intercept) | 927.63 | 282.48 | < 0.001 *** |
| Offspring diet | 4.13 | 1.26 | 0.265 |
| Maternal diet | 4.27 | 1.30 | 0.257 |
| Paternal diet | 33.05 | 10.06 | < 0.001 *** |
| Offspring diet : maternal diet | 5.41 | 1.65 | 0.202 |
| Offspring diet : paternal diet | -19.50 | 266.53 | 0.649 |
| Offspring diet : maternal diet :paternal diet[†] | 52.70 | 8.02 | < 0.001 *** |

† df=2, all other *df* =1, df res=103

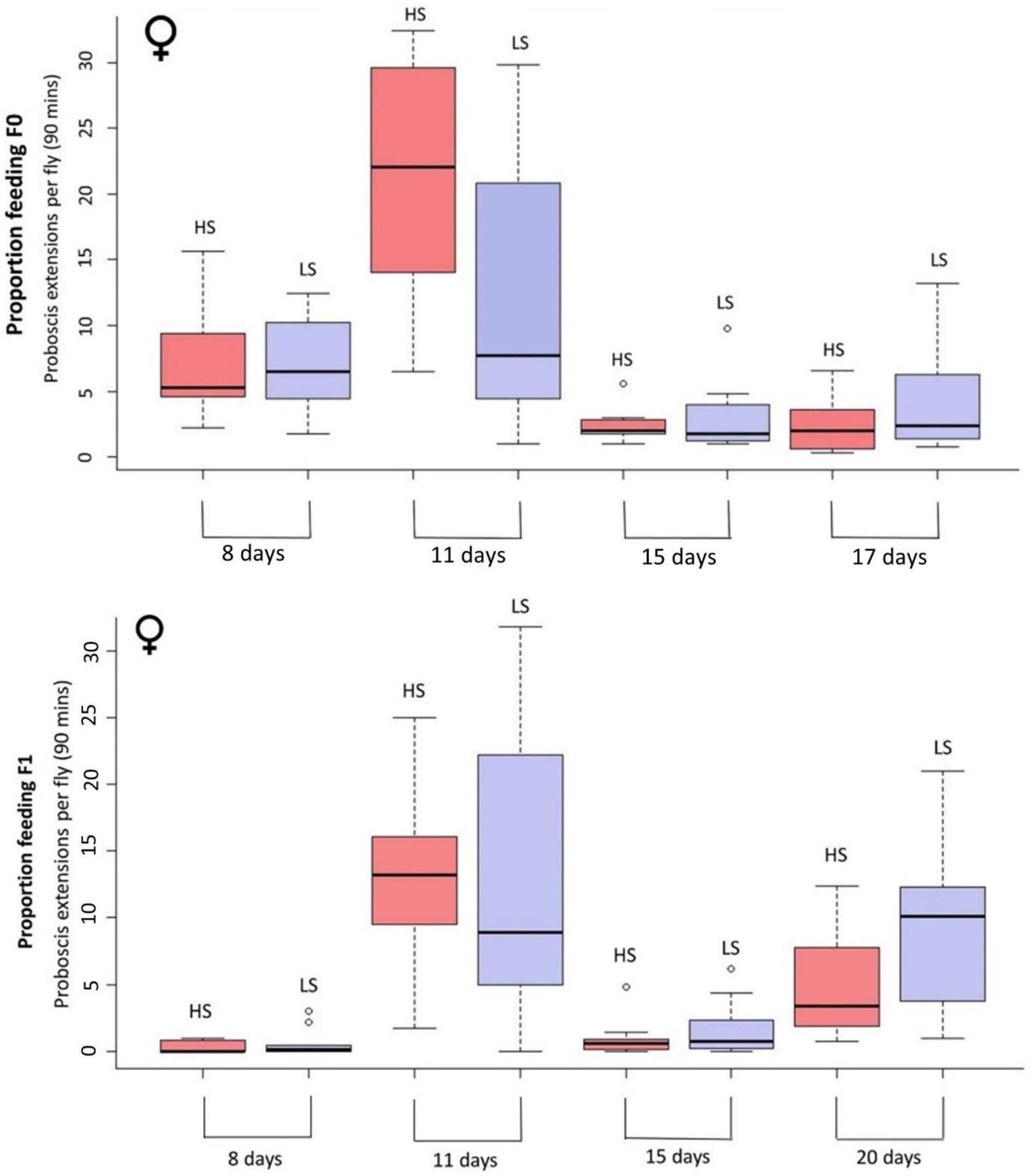


**Figure S1.** Median and quartiles of observed feeding for both generations, across 4 times periods. This is the total number of proboscis extensions into food media within 90 minutes, for the males and females of the parental generation at each age assayed. HS indicates a high sucrose diet of 20% (P:C ratio 1:5.3), LS indicates a low sucrose diet of 2.5% (P:C ratio 1:1.4)

### F1 TAG

**Table S9. A**nalysis of deviance table with Kenwood-Rogers F test, linear mixed model (Gaussian error distribution) after model reduction for predictors of offspring whole-body triglycerides.

*** indicates significance at $p < 0.001$, ** indicates significance at $p < 0.01$, * indicates significance at $p < 0.05$.

| *Fixed Effects* | | *F* | *p-value* |
|---|---|---|---|
| Diet | | 26.73 | < 0.001 *** |
| Sex | | 4.09 | < 0.05 * |
| Maternal diet | | 2.76 | 0.106 |
| Paternal diet | | 1.22 | 0.278 |
| Diet : sex | | 1.845 | 0.183 |
| Sex : maternal diet | | 3.49 | 0.070 |
| Sex : paternal diet | | 0.243 | 0.625 |
| Diet : maternal diet | | 23.78 | < 0.001 *** |
| Diet : paternal diet | | 2.23 | 0.144 |
| Diet : maternal diet : paternal diet[a] | | 4.28 | < 0.01 ** |
| Sex : maternal diet : paternal diet | | 1.19 | 0.283 |
| Diet : sex: maternal diet : paternal diet[b] | | 3.96 | < 0.01 ** |
| ***Random Effects*** | ***Variance*** | | |
| Vial identification | 0.000 | | |
| Residual | 0.011 | | |

[a]*df* = 2; [a]*df* = 3; all other *df* =1, df res=32

#### F1 Body mass

**Table S10.** Statistical results (analysis of deviance) of the linear mixed model (Gaussian error distribution) after model reduction for predictors of offspring weight (milligrams).
*** indicates significance at $p < 0.001$, ** indicates significance at $p < 0.01$, * indicates significance at $p < 0.05$.

| *Fixed Effects* | | *F* | *p-value* |
|---|---|---|---|
| (Intercept) | | 10560.11 | < 0.001 *** |
| Offspring diet | | 0.044 | 0.947 |
| Offspring sex | | 3936.54 | < 0.001 *** |
| Maternal diet | | 1.50 | 0.224 |
| Paternal diet | | 1.30 | 0.256 |
| Maternal diet : paternal diet | | 0.8921 | 0.347 |
| Offspring diet : maternal diet | | 7.08 | < 0.01 ** |
| Offspring diet : mat diet : pat diet | | 4.50[a] | < 0.05 * |
| ***Random Effects*** | ***Variance*** | | |
| Vial identification | 0.001 | | |
| Residual | 0.006 | | |

a)df=2, all other *df* =1, df res=86

#### F1 Feeding behaviour

**Table S11.** Statistical results (analysis of deviance with Kenwood-Roger's method) of the linear mixed model (Gaussian error distribution) after model reduction for predictors of offspring feeding behaviour.
*** indicates significance at $p < 0.001$, ** indicates significance at $p < 0.01$, * indicates significance at $p < 0.05$.

| ***Fixed Effects*** | | ***F*** | ***p-value*** |
|---|---|---|---|
| Diet | | 4.89 | 0.54 |
| Sex | | 24.85 | < 0.001 *** |
| Maternal diet | | 5.57 | 0.89 |
| Paternal diet | | 24.45 | 0.38 |
| Age | | 2.95 | 0.12 |
| ***Random Effects*** | ***Variance*** | | |
| Vial identification | 0.000 | | |
| Residual | 5.5 | | |

All *df* =1, df res=43

##### Correlations between body composition and fitness

##### Female F1 offspring

There is a weak negative relationship between female offspring triglyceride content and the amount of eggs they output, (r = -0.152, p = 0.718, BCa CI: -0.9519, 0.4283), and a moderate positive relationship between their triglyceride content and their lifespan (r = -0.399, p = 0.328, BCa CI: -0.5317, 0.9062). The relationship between female offspring body mass and egg output is a weak positive correlation (r = 0.214, p = 0.610, BCa CI: -0.6457, 0.8833), so too for female offspring body mass and lifespan, a weak positive correlation (r = 0.180, p = 0.668, BCa CI: -0.6298, 0.7948).

##### Male F1 offspring

For male offspring, a weak positive correlation was found between their triglyceride content and their lifespan (r = 0.196, p = 0.641, BCa CI: -0.7640, 0.8620), whereas a weak negative relationship was found between their body mass and their lifespan (r = -0.301, p = 0.469, BCa CI: -0.8132, 0.3609).

#### Further diet information

**Table S12**

| Sucrose | S:Y | P:C:F |
|---|---|---|
| 0.25% | 1:40 | 1 : 0.9 : 0.02 |
| 2.5% | 1:4 | 1 : 1.4 : 0.02 |
| 5% | 1:2 | 1 : 1.9 : 0.02 |
| 10% | 1:1 | 1 : 3.1 : 0.02 |
| 20% | 2:1 | 1 : 5.3 : 0.02 |
| 40% | 4:1 | 1 : 9.8 : 0.02 |

Calculated by overall mass of ingredients and not calories. (If calculating by calories the kcals of protein and carb are the same 4kcal/g and fat yields about 9ckal/g)

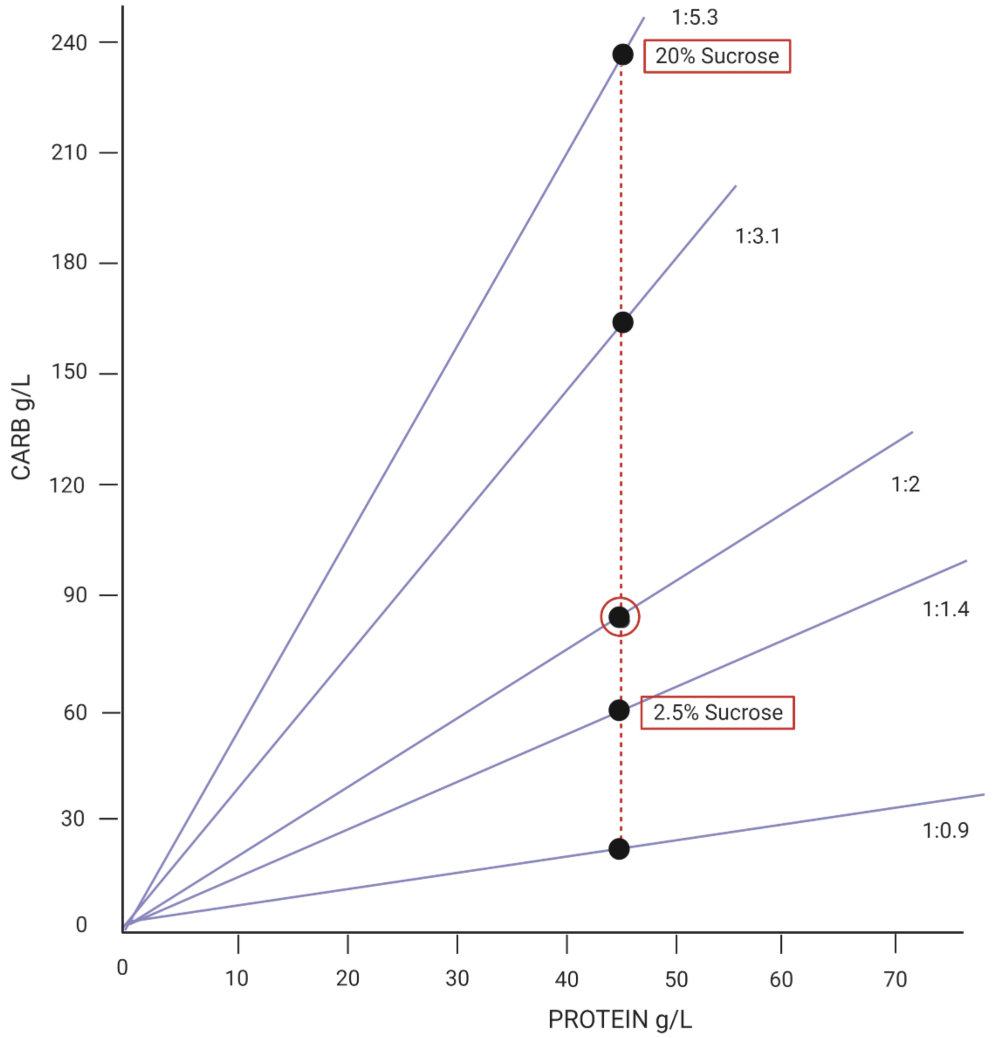

**Figure S2.** Sucrose levels in nutritional space

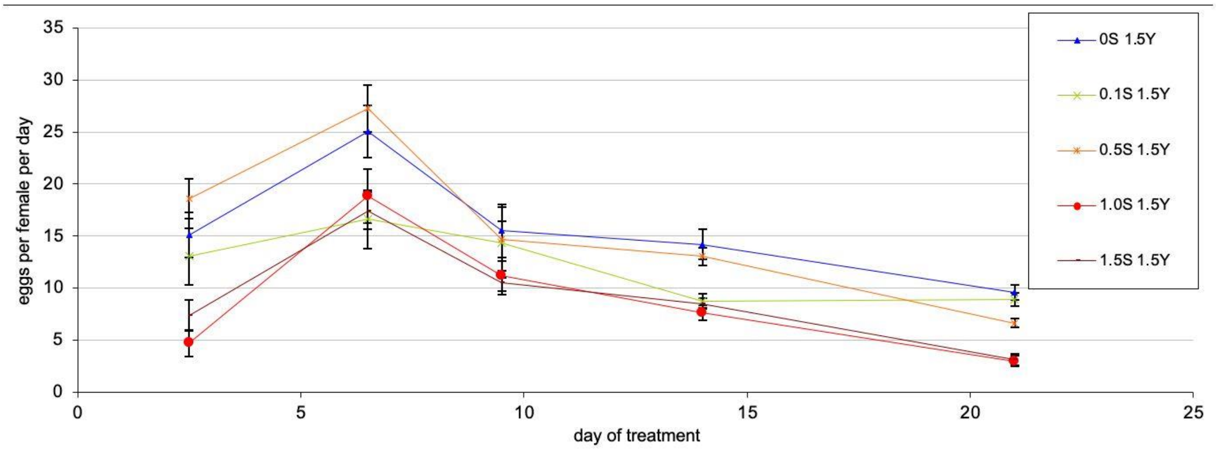


**Figure S3.** Reproductive peaks, and time points, under differing sucrose concentrations, with data from Bass et al., 2007.

**Lipid and protein Assay protocol**

Frozen flies (stored at -80C) were homogenized, using 5 flies per Eppendorf tube. Using a pestle, flies frozen in liquid nitrogen were crushed and 1 ml 0.05% Tween 20 lysis solution was added. Microtubes were then vortexed for 20 seconds each. We heated samples at 70°C in a waterbath for 5 min, and centrifuged at 5,000 rpm for 1 minute. We transferred supernatant (500 µl) from each sample to new Eppendorf tubes and centrifuged at 14,000 rpm for 3 minutes. Supernatant (50 µl) from each sample was then transferred to a 96-well plate, and 200 µl of Thermo Infinity Triglyceride solution (pre-warmed at 37°C) was added to each well. Samples were incubated for 5 minutes, and absorbance was measured at 540 nm.
